# A co-opted ISG15-USP18 binding mechanism normally reserved for deISGylation controls type I IFN signalling

**DOI:** 10.1101/2021.06.01.446527

**Authors:** Andri Vasou, Katie Nightingale, Vladimíra Cetkovská, Jonathan Scheler, Connor G.G. Bamford, Jelena Andrejeva, Ulrich Schwarz-Linek, Richard E. Randall, John McLauchlan, Michael P. Weekes, Dusan Bogunovic, David J Hughes

**Author notes:** Bioarchitech Ltd, Oxford UK (AV), School of Biological Sciences & Institute for Global Food Security, Queen’s University, Belfast UK (CGGB).

## Abstract

Type I interferon (IFN) signalling induces the expression of several hundred IFN-stimulated genes (ISGs) that provide an unfavourable environment for viral replication. To prevent an overexuberant response and autoinflammatory disease, IFN signalling requires tight control. One critical regulator is the ubiquitin-like protein ISG15, evidenced by autoinflammatory disease in patients with inherited ISG15 deficiencies. Current models suggest that ISG15 stabilises USP18, a well-established negative regulator of IFN signalling. USP18 also functions as an ISG15-specific peptidase that cleaves ISG15 from ISGylated proteins; however, USP18’s catalytic activity is dispensable for controlling IFN signalling. Here, we show that the ISG15-dependent stabilisation of USP18 involves transient hydrophobic interactions. Nonetheless, while USP18 stabilisation is necessary, it is not sufficient for regulation of IFN signalling. USP18 requires non-covalent interactions with the ISG15 C-terminal diGlycine motif to promote its regulatory function. This trait may have been acquired in humans through co-option of a binding mechanism normally reserved for deISGylation, identifying an unexpected new function for human ISG15.

## Introduction

The interferon (IFN) response plays a critical role in orchestrating protective antiviral immune responses to combat viral infections ^1^. Type I IFNs are widely expressed by all nucleated cells following viral infection ^2^ and signal through the IFN-α/β receptor (IFNAR), which consists of subunits IFNAR1 and IFNAR2 ^3^. Engagement of IFNAR triggers a phosphorylation cascade involving the reciprocal trans-phosphorylation of Janus kinase 1 (JAK1) and tyrosine kinase 2 (Tyk2) ^4^ and phosphorylation of the cytoplasmic tails of the receptor subunits creating a docking site for the recruitment and subsequent phosphorylation of signal transducer and activator of transcription 1 (STAT1) and STAT2 ^5^. Once activated, STAT1/2 heterodimers associate with IFN-regulatory factor 9 (IRF9) to form IFN-stimulated gene factor 3 (ISGF3), which binds to the IFN-stimulated response element (ISRE) within the promoters of interferon stimulated genes (ISGs) ^1^. Several ISGs have been shown to have specific antiviral activity and/or play a role in regulating the IFN response itself. Although this is a prompt and powerful defence against pathogens, a dysregulated type I IFN response can lead to autoinflammatory disease. Therefore, tight regulation of activating and inhibitory signals is of paramount importance for maintaining the protective host-defence nature of the response but limiting tissue damage.

ISG15, a ubiquitin-like modifier, is synthesised from a precursor that is cleaved at the C-terminus to yield the mature 15-kDa protein with a C-terminal Leu-Arg-Leu-Arg-Gly-Gly (LRLRGG) tail ^6,7^. ISG15 exists as a free molecule but can also covalently bind to target proteins through the formation of an isopeptide bond between its terminal glycine and the lysine ε-amino group of the target protein ^8^, a process termed ISGylation (reviewed by ^9,10^). ISGylation is reversible through the action of a deISGylase enzyme, the ubiquitin-specific protease 18 (USP18) ^11^. As a protease, USP18 specifically deconjugates ISG15 from modified proteins and shows no reactivity towards other ubiquitin-like proteins. This specificity is achieved through hydrophobic interactions centred on Trp121 (W121) in mouse Isg15 (Trp123 in humans) ^12^. Independent to its isopeptidase activity on ISG15 ^13,14^, USP18 interacts with STAT2 to facilitate its recruitment to IFNAR2, where it can inhibit receptor dimerization by interfering with cytoplasmic interactions between IFNAR subunits ^15–17^.

Several reports now show that inherited ISG15-deficiency in humans causes type I interferonopathy and autoinflammatory disease ^18–20^. ISG15-deficient cells exhibited enhanced and prolonged ISG expression and a concomitant resistance to virus infection ^19,21,22^ a phenotype also associated with USP18 deficiency ^14,23^. Indeed, despite high levels of *USP18* transcription, USP18 protein levels in ISG15-deficient cells are very low ^19,21,22^ and previous reports have shown that intracellular ISG15 is required for rescuing USP18 from S-phase kinase associated protein 2 (SKP2)-mediated proteasomal degradation ^19,24^.

Here, contrary to existing models, we show that the ISG15-dependent stabilisation of USP18 is necessary *but not sufficient* for regulation of type I IFN signalling. ISG15-dependent stabilisation of USP18 requires hydrophobic interactions coordinated by ISG15-Trp123, but this interaction is weak and is likely regulated by the relative abundance of ISG15 and USP18. Nevertheless, independently of ISG15’s ability to stabilise USP18, we show that high affinity ISG15-USP18 interaction via the ISG15 C-terminal di-Gly motif is required to negatively regulate type I IFN signalling, and abolishing this non-covalent interaction results in phenotypes associated with enhanced IFN-α signalling. Together our data illustrate that a binding mechanism normally reserved for deISGylation has likely been co-opted to serve a crucial role in regulating early intracellular immune responses in humans.

## Results

### The C-terminal di-Gly motif of ISG15 is important for IFN-α signalling regulation

In work leading to this report, we found that reconstituting expression of C-terminal mutants of ISG15, where the terminal di-Gly motif was replaced with di-Ala, in ISG15-deficient cells did not restore the regulation of IFN signalling. To dissect the role of the ISG15 C-terminus, we reconstituted expression of Myc-tagged ISG15 (using lentiviral transduction), which retained its di-Gly motif (ISG15.GG), or C-terminal mutants of ISG15, where the di-Gly motif was either replaced by di-Ala (ISG15.AA) or deleted (ISG15.ΔGG), in our phenotypically validated A549-ISG15^−/−^ cell line ^22^. To better mimic physiological conditions, reconstituted ISG15 was placed under the control of the native ISG15 promoter (pr15) (Fig. 1A). IFN-α stimulation induced expression of Myc-ISG15 to levels similar to endogenous ISG15 in control A549s and, as expected, ISG15.ΔGG and ISG15.AA did not ISGylate proteins (Fig. 1B). These experimentally tractable cells are therefore valuable for deciphering the role of ISG15 or other ISGs during the innate immune response.

**Figure 1.**
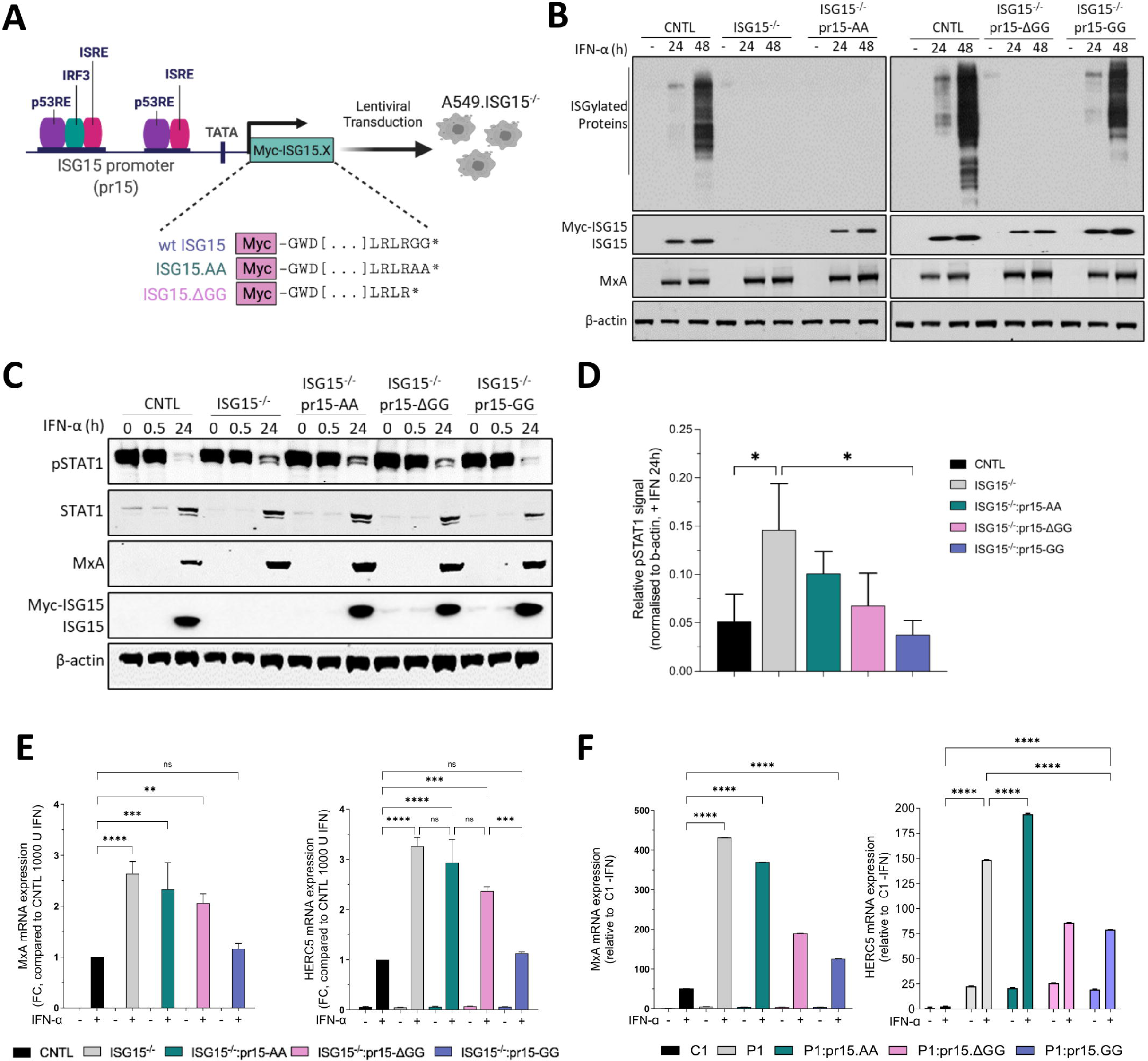
Functional characterisation of A549-ISG15^−/−^ cell lines reconstituted with the C-terminal ISG15 mutants. **(A)** Schematic presentation of the lentiviral technology used to reconstitute ISG15 expression in A549-ISG15^−/−^ using an inducible system where expression of ISG15 is under the control of its native promoter (pr15). The induction of pr15 is regulated by transcription factors physiologically upregulated by innate immune responses (e.g., IRF3 and ISGF3) or cell-cycle regulators (e.g., p53) **(B)** Immunoblot analysis of ISG15 expression induced by IFN-α treatment. A549 (CNTL), ISG15^−/−^ and ISG15.AA-, ΔGG-, GG-expressing derivatives were treated with 1000 IU/mL IFN-α for 24 and 48 h or left untreated. Whole cell extracts were prepared and ISG15, MxA and β-actin proteins were analysed by immunoblot. **(C)** A549 (CNTL), 1SG15^−/−^ and ISG15.AA-, ΔGG-, GG-expressing derivatives were treated with 1000 IU/mL IFN-α for 30 min, then extensively washed and media without IFN replaced. Cells were harvested at 0 and 30 min and 24 h after IFN-α removal and phopho-STAT1, total STAT1, MxA, ISG15 and β-actin were detected by immunoblot. **(D)** Experiments in (C) were performed on three independent occasions and phospho-STAT1 levels after 24 h IFN-α removal were quantified using Image Studio software (LI-COR). Error bars represent the SD of the mean and statistical significance was assessed using one-way ANOVA and Sidak post-tests. Data in lanes 1 – 6 have previously been reported (21) **(E)** A549 (CNTL), ISG15^−/−^ and ISG15.AA-, ΔGG-, GG-expressing derivatives were treated with 1000 IU/mL IFN-α for 24 h. Expression of ISGs was tested using reverse transcription quantitative PCR (RT-qPCR) with primers specific for MxA and HERC5. Relative expression was determined following SYBR Green quantitative PCR (qPCR) using ΔΔCt method. β-Actin expression was used to normalize between samples. Data are presented as a mean fold increase relative to IFN-α-treated A549 control cells (set to 1). **(F)** hTert-immortalized ISG15-deficient patient-derived dermal fibroblasts (P1) transduced with ISG15.GG (wt) and C-terminal mutants (ISG15.AA and ISG15.ΔGG) were treated with 1000 IU/ml IFN-α for 12 h, or left untreated, washed and incubated in media without IFN-α for 24 h. MxA and HERC5 expression was analysed as in (E) with data presented as fold increase relative to non-IFN-α treated P1. Error bars (in E and F) represent the SD of the mean from three independent experiments performed on different occasions. Each experiment additionally included three technical replicates. Statistical significance was assessed using two-way ANOVA and Tukey multiple comparisons test; *, p < 0.05, **, p < 0.01, ***, p < 0.001, ****, p < 0.0001, n.s., no statistical significance.

We have previously shown that IFN-α treatment of A549-ISG15^−/−^ cells results in enhanced signalling characterised by elevated phospho-STAT1 levels and an augmented and prolonged expression of ISGs ^22^. Here we determined the ability of C-terminal mutants of ISG15 to regulates signalling. Cells were treated with IFN-α for 30 min, extensively washed, and re-incubated in media without IFN-α. Cell lysates taken immediately after 30-min treatment (0 min) and 30 min later (0.5 h) showed high levels of STAT1 phosphorylation in all tested cell lines (Fig. 1C). Following 24-h treatment with IFN-α, there was evident expression of the ISGs MxA and ISG15, Myc-ISG15 and enhanced expression of STAT1 (Fig. 1C). Interestingly, although phospho-STAT1 levels had declined in A549 control and ISG15.GG-expressing cells at 24 h after IFN-α treatment, levels were clearly higher in A549-ISG15^−/−^ cells and the cells expressing ISG15.AA and ISG15.ΔGG, signifying higher levels of IFN-α signalling in these cells (Fig. 1D).

To confirm these observations at the level of ISG transcripts, cells were treated with IFN-α for 24 h or left untreated. As expected, expression of *MxA* and *HERC5* mRNA was significantly higher in A549-ISG15^−/−^ cells compared to A549 control (2.6- and 3.3-fold respectively, Fig. 1E). Consistent with the phosho-STAT1 levels, ISG mRNA levels were also significantly higher in cells expressing the ISG15.AA and ISG15.ΔGG mutants. Intriguingly, this demonstrated a gradient pattern of regulation where ISG15.GG-expressing cells regulated similar to control A549 cells, cells expressing the ISG15.AA mutant were characterised by higher ISG levels similar levels to A549-ISG15^−/−^ cells, but cells expressing the ISG15.ΔGG mutant consistently showed a trend towards an intermediate level of regulation. No significant difference was observed between the ISG15.GG-expressing cells and the control, suggesting that the level of ISG15 expression in this system is sufficient to regulate IFN signalling similarly to unmodified control cells (Fig. 1E). We also transduced hTert-immortalized skin fibroblasts from ISG15^−/−^ patient cells (P1) with the same ISG15 vectors as the A549 model and investigated ISG expression compared to hTert-immortalized skin fibroblasts from a ISG15^+/+^ donor (C1) following treatment with IFNα (Fig. 1F). ISG expression (*MxA* and *HERC5*) was higher in P1 and all reconstituted P1 cells than C1 cells regardless of IFN treatment which possibly reflects that, unlike the isogenic A549 cells, C1 and P1 are from different donors (hence, what complete regulation looks like in these patients cannot be known). Nevertheless, the profile of ISG expression, where expression was highest in ISG15-deficient cells followed by ISG15-AA, then -ΔGG, then -GG, was also observed in the patient cells. Overall, these data show that the C-terminal di-Gly motif is important for the negative regulation of type I IFN signalling.

### IFN-α-pretreatment leads to viral resistance in cell lines expressing the C-terminal mutants of ISG15

We and others have previously shown that pre-treatment of ISG15-deficient cells with IFN-α renders them resistant to viral infection ^21,22^. Because we have reported that viral resistance in ISG15-deficient cells is directly related to dysregulated type I IFN signalling ^22^, this assay serves as an excellent model for investigating the biological implications of ISG15 loss-of-function and the regulatory role of the ISG15 di-Gly motif. Cell lines were primed with IFN-α for 18 h, then infected with a recombinant parainfluenza virus type 5 (PIV5) expressing mCherry (rPIV5-mCherry). Because the PIV5 V-protein targets STAT1 for proteasomal degradation, rPIV5-mCherry can replicate in A549 control cells despite a primed IFN response, albeit with reduced kinetics ^25^. By 48 h of infection in A549 control cells, there was a corresponding increase in PIV5 NP expression, STAT1 was undetectable and MxA expression was reduced (Fig. 2A). As anticipated, pretreatment with IFN-α rendered A549-ISG15^−/−^ cells resistant to PIV5 infection, whereas reconstituted expression of ISG15.GG reversed the phenotype (Fig. 2A,B). Interestingly, like A549-ISG15^−/−^ cells, NP expression was undetectable at 24 h post-infection in both ISG15.AA and ISG15.ΔGG-expressing cells (Fig. 2A). Although PIV5 infection recovered to some extent by 48 h, NP abundance was still significantly reduced by 90% and 45% at 48 h p.i. in cells expressing ISG15.AA and ISG15.ΔGG, respectively (Fig. 2B). Intriguingly, this gradient in viral protein expression inversely correlated with the gradient observed for ISG expression (Fig. 1E), linking the ability of ISG15 and ISG15 mutants to regulate the magnitude of the antiviral response with their capacity to support infection. Moreover, we used fluorescence microscopy to visualise mCherry expression and indicate rPIV5-mCherry replication. Consistent with NP expression, no mCherry was detected in the A549-ISG15^−/−^ cells (Fig. 2C) or ISG15-deficient patient cells (P1) (Fig. 2D), with low or moderate levels in cells expressing the ISG15.AA and ISG15.ΔGG mutants, respectively (Fig. 2C,D). Altogether, these experiments highlight the impact on viral infection of the C-terminal di-Gly motif of ISG15, via its regulation of IFN signalling.

**Figure 2.**
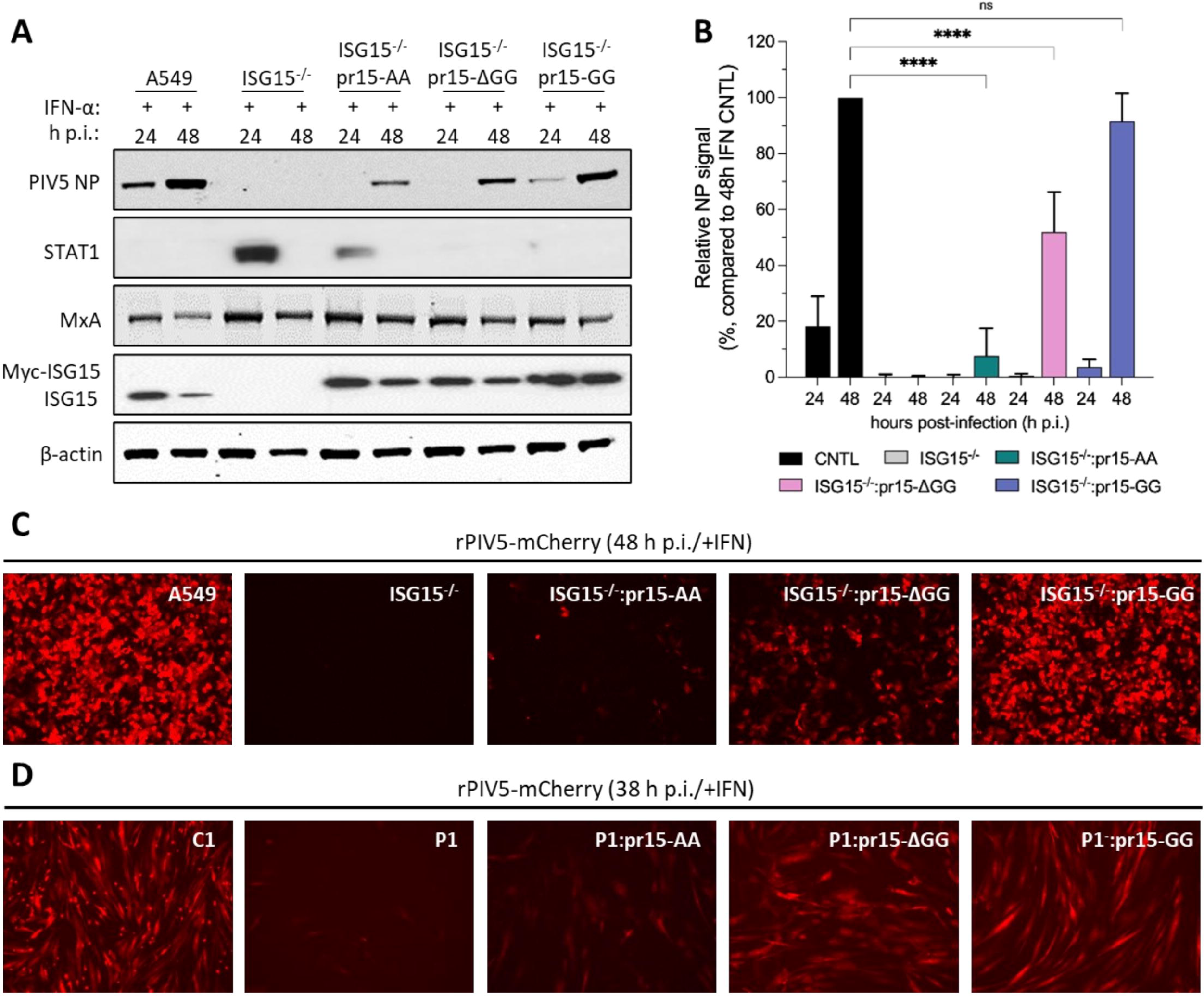
The ability of ISG15 C-terminus to regulate IFN-α signalling influences viral infection. **(A)** A549 (CNTL), ISG15^−/−^ and ISG15.AA-, ISG15.ΔGG- and ISG15.GG-expressing derivatives were treated with 1000 IU/mL IFN-α for 18 h and then infected with rPIV5-cherry (MOI 10). Cells were harvested at 24 and 48 h p.i. and processed for immunoblot analysis using antibodies specific for PIV5 NP, STAT1, MxA, ISG15 and β-actin. Reconstituted ISG15 expression was under the control of a cloned ISG15 promoter (pr15) and was therefore IFN-inducible. **(B)** Experiments described in (A) were performed independently three times (infections were performed on three separate occasions), and NP and β-actin levels were quantified using Image Studio software (LI-COR Biosciences). Signals were relative to those generated from IFN-α-treated A549 cells infected for 48 h p.i. (set to 100%). Error bars = SD. Statistical significance was assessed using two-way ANOVA and Tukey multiple comparisons test; ****, p < 0.0001, n.s., no statistical significance. **(C)** Fluorescent imaging of mCherry expression, indicative of rPIV5-mCherry infection in A549, ISG15^−/−^ and ISG15.AA-, ISG15.ΔGG- and ISG15.GG-expressing A549-ISG15^−/−^. Cells were treated with 1000 IU/mL IFN-α for 18 h and then infected with rPIV5-mCherry (MOI 10) and imaged at 48 h post infection (h.p.i.). **(D)** Experiments described in (C) were performed with hTert-immortilized dermal fibroblasts. C1 are cells from ISG15^+/+^ donor and P1 are from an ISG15^−/−^ donor. Images were taken 38 h p.i.

### The ISG15-dependent stabilisation of USP18 is necessary but not sufficient for the regulation of IFN signalling

It has been established that ISG15 is crucial for sustaining the levels of USP18, a key negative regulator of IFN signalling, by preventing its SKP2-mediated ubiquitination and proteasomal degradation ^19,21,24^. Therefore, we reasoned that modifications to the ISG15 C-terminus might have affected its ability to stabilise USP18. To test this, cells were treated with IFN-α for 24 and 48 h and cell lysates were subjected to immunoblot analysis. Remarkably, although USP18 abundance was dramatically reduced in A549-ISG15^−/−^ cells, its levels in ISG15.AA and ISG15.ΔGG expressing cells were comparable to those observed in ISG15.GG-expressing and control A549 cells (Fig. 3A).

**Figure 3.**
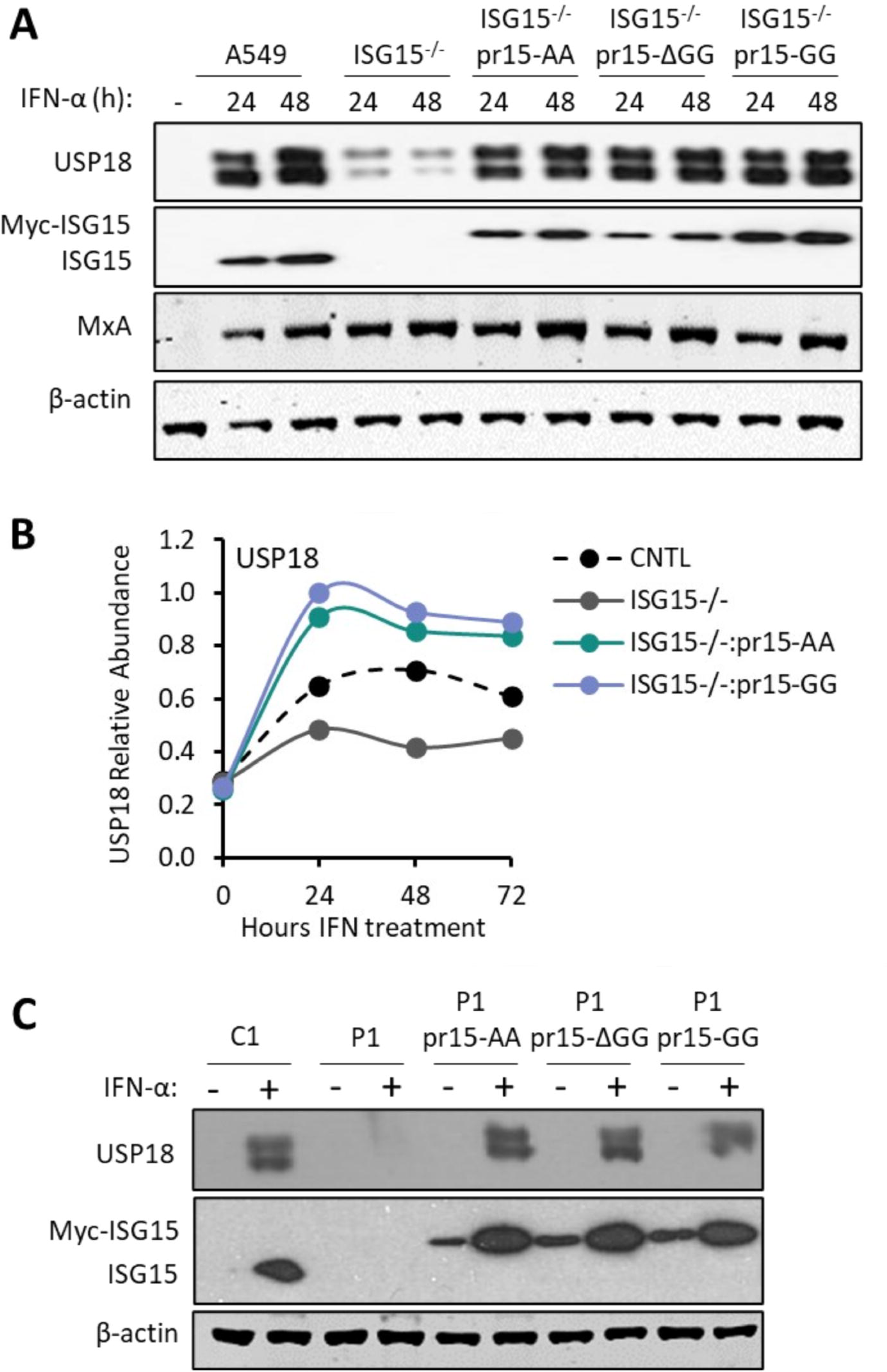
The C-terminal di-Gly motif of ISG15 is not required for USP18 stabilisation. **(A)** Immunoblot analysis of USP18 expression in A549 (CNTL), ISG15^−/−^ and ISG15.AA-, ΔGG-, GG-expressing derivatives after treatment with 1000 IU/mL IFN-α for 24 h. Whole cell extracts were prepared and USP18, ISG15, MxA and β-actin protein levels were analysed by immunoblot. **(B)** USP18 abundance in A549 (CNTL), ISG15^−/−^ and ISG15.AA and ISG15.GG-expressing derivative cells measured using quantitative tandem mass tags (TMT)-based proteomic analysis. Indicated cells were treated with 1000 IU/mL IFN-α for the indicated times (‘0 h’ cells were not treated with IFN-α and harvested at 72 h). **(C)** hTert-immortilized dermal fibroblasts from a ISG15^+/+^ donor (C1), from an ISG15^−/−^ donor (P1) and P1 cells reconstituted with IFN-inducible (pr15) ISG15.AA, ISG15.ΔGG and ISG15.GG. Cells were treated with 1000 IU/mL IFN-α, or not treated, for 48 h. Whole cell extracts were prepared and USP18, ISG15 and β-actin protein levels were analysed by immunoblot.

Next, we employed a multiplexed proteomic approach to obtain an unbiased, global picture of the proteomic changes induced by IFN-α and to independently assess the ability of ISG15 to stabilise USP18. A549 control cells, A549-ISG15^−/−^ cells and knockout cells expressing IFN-inducible ISG15.AA or ISG15.GG were treated with IFN-α for 24, 48 or 72 h, and whole cell protein abundance measured using 16-plex TMT labelling and MS3 mass spectrometry (Supplementary Fig. 1). In total, 7112 proteins were quantified (full data can be visualised using the ‘Plotter’ in Supplementary Table 1). Importantly, the proteomics analysis provided verification that ISG15 expression, independently of its ability to regulate signalling, was able to stabilise USP18 (Fig. 3B). We also verified that USP18 was stabilised in ISG15-deficient patient cells expressing all forms of ISG15 (Fig. 3C).

Further analysis of the proteomics data showed that, consistent with gene expression analyses, the abundance of IFN-stimulated proteins, such as ISG20 and IFIT1 was higher in the A549-ISG15^−/−^ and ISG15.AA-expressing cells compared to the A549 control and ISG15.GG-expressing cells (Supplementary Fig. 1B). To determine whether this trend was reflected more generally, ISGs were identified by (a) comparison to the Interferome database ^26^, and (b) stringent criteria to identify proteins upregulated by IFN-α compared to mock treatment in A549 control cells (>1.7-fold increase in abundance). Ninety proteins identified by these criteria as interferon stimulated were expressed at significantly higher levels by A549-ISG15^−/−^ and ISG15.AA cells compared to both A549 control and ISG15.GG cells (Supplementary Fig. 1C). As expected, there was no statistically significant difference in ISG expression between A549 control and ISG15.GG-expressing cells, providing further evidence that genetic manipulation did not cause any inadvertent phenotypic changes to the cells.

To determine whether specific pathways were enriched in the ISG15^−/−^ and ISG15.AA cells, we used DAVID analysis to examine proteins upregulated >1.7-fold compared to control A549 cells. As anticipated, effector molecules involved in cellular immune responses, such as innate immunity and antiviral defence, were enriched in all tested cell lines with several overlapping factors in each cluster (Supplementary Fig. 1D). Although fewer pathways were enriched in ISG15.GG cells suggesting tighter control than in cells expressing C-terminal mutants, antiviral defence, immunity and innate immunity pathways were identified suggesting that their control was not as tightly regulated compared to control A549 cells. This might be because, despite being inducible by IFN-α, lentivirally-delivered ISG15 transgenes are not in their authentic genomic loci, or that lack of splicing due to expression of a cDNA, affects expression. Interestingly, components of IFN-γ mediated signalling and antigen processing and presentation pathways were selectively enriched in A549-ISG15^−/−^ and ISG15.AA-expressing cells, for example transporter associated with antigen presentation 1 (TAP1) (Supplementary Fig. 1D-E). One potential explanation is the selective enrichment in these cells of key regulatory factors induced by primary IFN-γ signalling ^27,28^, such as the interferon-regulatory factor 1 (IRF1) whose expression is induced by STAT1 homodimers typical of IFN-γ-induced ISGs with gamma-activated sequences (GAS) in their promoters and not STAT1-STAT2-IRF9 (ISGF3) complexes that characteristically induce genes with IFN-stimulated response elements (ISRE) activated by type I IFN signalling (although GAS genes can be activated upon type I IFN signalling as concentrations of STAT1-homodimers increase; Supplementary Fig. 1E). It may be that enrichment with components of IFN-γ signalling derives through the enhanced levels of active STAT1 (Fig. 1C) and subsequent increased stoichiometry of STAT1 homodimer complexes in ISG15^−/−^ and ISG15.AA cells, leading to gene expression through binding to GAS elements in the promoters of ISGs ^29^. Moreover, cell cycle components and several factors with nucleotide binding properties were also enriched in cells with a dysregulated IFN-α signalling response (Supplementary Fig. 1D). In conclusion, using unbiased proteomic analyses, we have further confirmed that the C-terminal di-Gly domain of ISG15 is important for regulation of type I IFN responses. It is also possible that pathways associated with IFN-γ, which include genes associated with antiproliferative phenotypes and apoptosis, may underpin the pathogenesis of autoinflammatory diseases associated with loss of type I IFN signalling control. Moreover, these experiments clearly show that, although the stabilization of USP18 is crucial, it is not sufficient for the regulation of type I IFN signalling.

### Hydrophobic interactions are required for ISG15-dependent stabilization of USP18 but ISG15 di-Gly-dependent interactions are necessary for type I IFN signalling regulation

Because neither ISGylation ^19,22^ or ISG15-dependent stabilisation of USP18 are sufficient for the regulation of type I IFN signalling, ISG15’s function, with respect to the IFN pathway, is likely to involve a non-covalent protein-protein interaction. The most likely candidate is USP18, given that ISG15 and USP18 must interact during the deISGylation process and both their involvement in regulation. For the ISG15 C-terminal tail to engage with the catalytic pocket of USP18, a two-step binding process is required; initially, a hydrophobic patch in ISG15 (centred on Trp121 in mouse, Trp123 in human, Fig. 4A) must interact with a hydrophobic patch in USP18 (termed ISG15 binding box 1 (IBB-1) resulting in a rearrangement of the USP18 ‘switching loop’ thus allowing the ISG15 C-terminal tail access to the catalytic site in USP18. Importantly, these hydrophobic interactions explained the specificity of USP18 towards ISG15 and no other ubiquitin-like proteins ^12^. We modelled the hISG15-hUSP18 complex using AlphaFold2; even though the N-terminal 46 residues of USP18 were predicted to be intrinsically unstructured, AlphaFold2 predicted the ISG15:USP18 complex with very high confidence (>90) as measured using pLDDT or predicted local distance difference test ^30^ (Supplementary figure 2). These data showed that Ala141, Leu145, Pro195 and His255 in USP18 (equivalent to Ala138, Leu142, Pro195 and His251 in mUsp18) form the IBB-1 pocket that accommodates ISG15-Trp123 (Fig. 4A), which demonstrated a high degree of conservation between human and mouse complexes. We then mutated Trp123 to Arg in our IFN-regulated wt Myc-ISG15-GG vector, the equivalent mutation used by Basters et al. ^12^ to investigate the mouse complex, and reconstituted A549-ISG15^−/−^ cells. Upon stimulation with IFN-α, ISG15-W123R expressed to similar levels as endogenous ISG15 in control A549 cells (Fig. 4B). Interestingly, ISG15-W123R was able to ISGylate proteins showing this residue is not required for interactions with UBA7 (E1), UBE2L6 (E2) or HERC5 (E3) and, though not quantified, the degree of ISGylation appeared larger. However, USP18 was not stabilised in these cells suggesting that initial hydrophobic interactions are required to stabilise USP18. This observation also accounted for the observed increase in ISGylation, as in the absence of (or interaction with) USP18, ISG15 cannot be deconjugated and ISG expression, including the ISGylation machinery, would be increased. Indeed, *MxA* and *HERC5* expression, as examples of ISGs, was equivalent to expression in A549-ISG15^−/−^ and significantly higher than in control A549 cells (Fig. 4C). Next, we assessed the ability of endogenous USP18 to interact with ISG15 and its C-terminal mutants in our reconstituted cell lines. Myc-tagged ISG15 was immunoprecipitated from ISG15-reconstituted cells following a 24-h IFN-α treatment. To verify that ISGylation was not necessary for the interaction between ISG15 and USP18, we knocked-out UBA7 from our ISG15.GG-expressing cells by CRISPR/Cas9 genome editing (Fig. 4D). Here we showed that the ability of ISG15 mutants to bind USP18 mirrored the gradient pattern of ISG mRNA regulation (Fig. 1E) and corresponding effects on viral infection (Fig. 2). Reconstituted ISG15.GG efficiently bound USP18; however, the ISG15.AA mutant was unable to interact with USP18 (or its binding was below the limits of detection), whereas ISG15.ΔGG did interact but at reduced levels compared to ISG15.GG (Fig. 4D). Notably, USP18 co-immunoprecipitated with ISG15.GG in UBA7^−/−^ cells, confirming that the ISG15-USP18 interaction is not dependent on ISGylation (Fig. 4D). Furthermore, we noted that all forms of ISG15 stabilised USP18 (see WCL samples, Fig. 4D lower panels, though no ISG15^−/−^ control was included), again confirming that ISG15 stabilised USP18 independently of their ability to regulate signalling.

**Figure 4.**
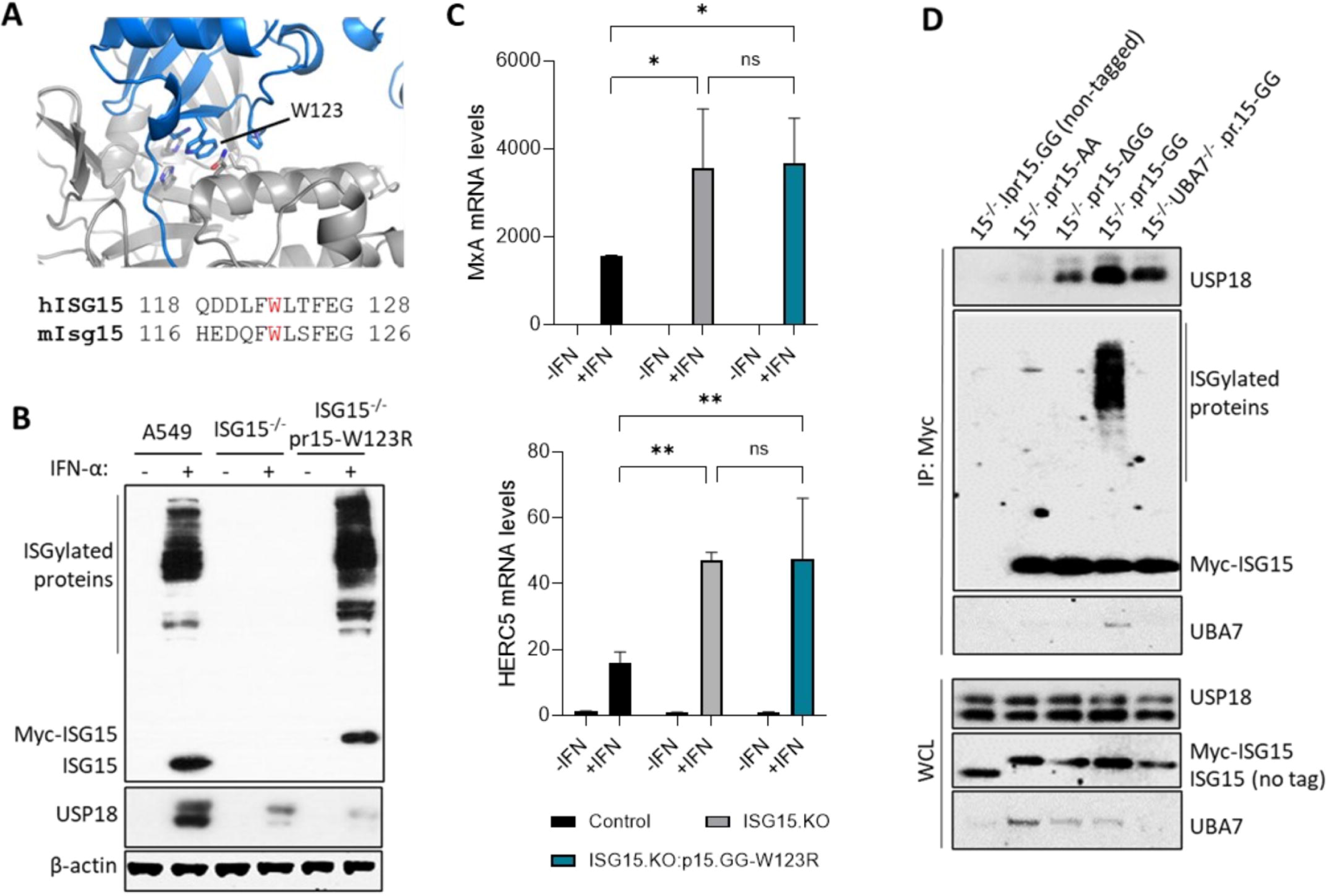
Tryptophan 123 (Trp123/W123) in ISG15 is required for USP18 stability and the C-terminal tail of ISG15 is crucial for the regulation of type I IFN signalling. **(A)** AlphaFold2 model representing a complex between the ISG15 binding box 1 (IBB-1) in hUSP18 (grey; Uniprot ID: Q9UMW8) and the hydrophobic pocket that accommodates the W123 side chain from hISG15 (blue; Uniprot ID: P05161) along with an amino acid sequence alignment of the corresponding region of mIsg15 (W121) and hISG15 (W123). **(B)** A549-ISG15^−/−^ were reconstituted with ISG15-W123R under the control of the ISG15 promoter (pr15) and was therefore IFN-inducible. A549, A549-ISG15^−/−^ and A549-ISG15^−/−^-pr15-W123R were treated with 1000 IU/mL IFN-α for 24 h. Whole cell extracts were prepared and USP18, ISG15 and β-actin protein levels were analysed by immunoblot. **(C)** A549, A549-ISG15^−/−^ and A549-ISG15^−/−^-pr15-W123R were treated with 1000 IU/mL IFN-α for 24 h and MxA and HERC5 expression was measured by RTqPCR (see Fig. 1E for details). **(D)** Immunoprecipitation of Myc-tagged ISG15.AA-, ΔGG- and GG after treatment with 1000 IU/mL IFN-α for 24 h. An ISG15^−/−^ cell-line expressing a non-tagged form of ISG15 was used as a negative control (left lane) and an ISG15.GG-expressing 1SG15^−/−^.UBA7^−/−^ cell line was used as an ISGylation-deficient control (final lane). ISG15 was immunoprecipitated (IP) using anti-c-Myc antibodies covalently coupled to magnetic beads. Immunoprecipitates (top) and whole cell lysates (WCL; bottom) were subject to immunoblot analysis with antibodies to USP18, ISG15 and UBA7. ISG15 expression in reconstituted cell lines was under the control of the ISG15 promoter (pr15) and was therefore inducible by IFN. Data representative of at least three independent assays.

Interestingly, that USP18 was stabilised in the presence of ISG15 C-terminal mutants suggests that ISG15 Trp123-USP18 IBB-1 interaction was intact; however, it was clear that the ISG15 C-terminal tail significantly contributed to affinity, as when this was mutated, any interaction was below the level of detection (ISG15.AA) or reduced (ISG15-ΔGG) compared to wild type ISG15 (Fig. 4D). Collectively, these results indicate that low affinity ISG15-W123-dependent interactions are required for USP18 stability but that the C-terminal di-Gly motif is important for enhanced binding and is necessary to facilitate USP18’s inhibition of type I IFN signalling.

### The ISG15-USP18 interaction is important for the tight regulation of IFN-α signalling

To independently assess the requirement of the ISG15-USP18 interaction for the regulation of IFN signalling, we mutated the USP18 isoleucine residue at position 60 (USP18.I60N), which is known to abolish interactions with ISG15 ^24^. In addition, since previous studies have shown that the mouse Usp18 (Ubp43) negatively regulates IFN signalling in the absence of its isopeptidase activity ^14^, we additionally investigated the catalytically inactive human mutant USP18.C64S. We first sought to evaluate the impact of these point-mutations on ISG15-USP18 binding by coimmunoprecipitation (co-IP) (Fig. 5A). Consistent with previous studies ^24^, our data demonstrated that the USP18.I60N mutant was unable to interact with ISG15.GG, whereas the USP18.C64S mutant had stronger binding affinity for ISG15 compared to wt USP18 ^24^ (Fig. 5A).

**Figure 5.**
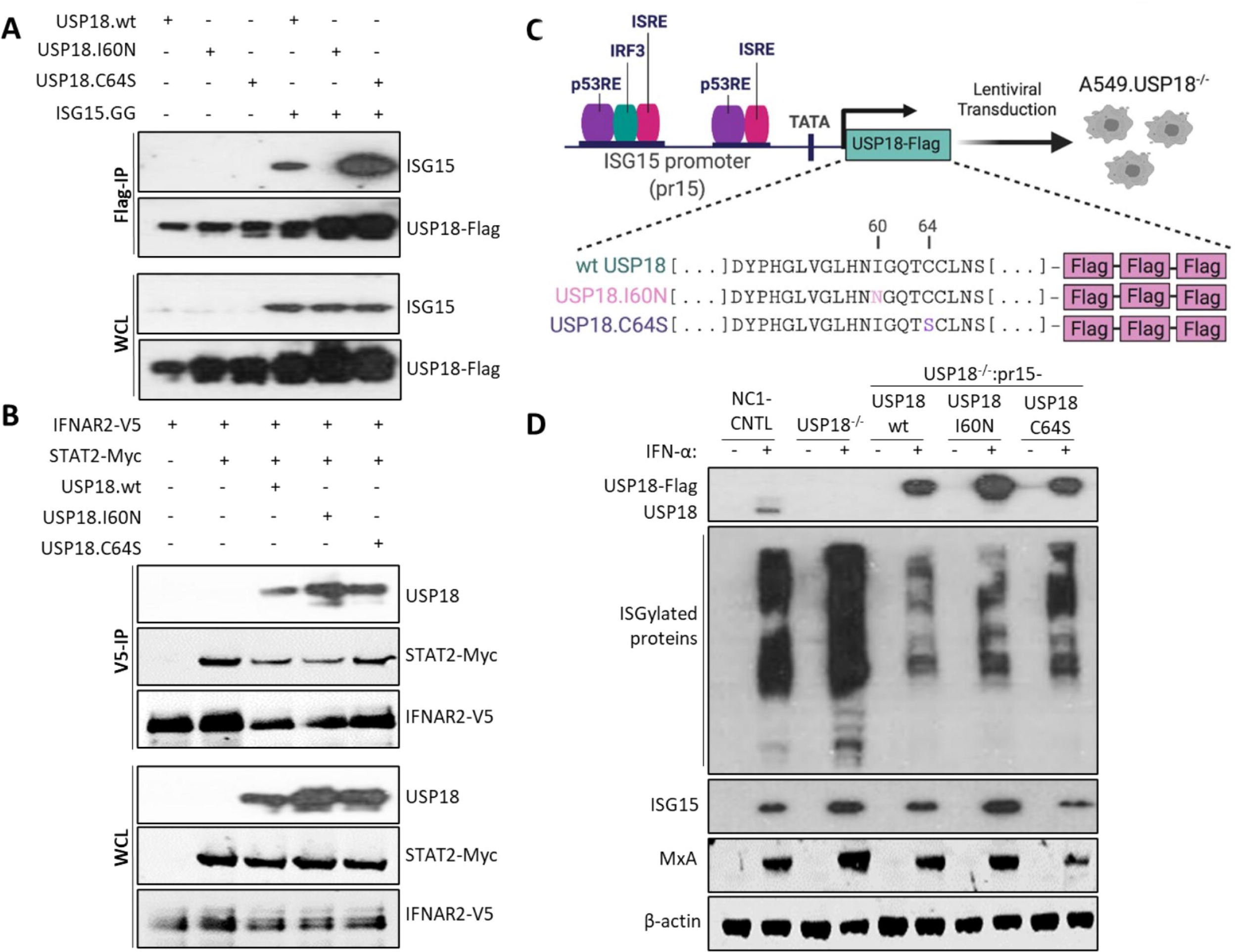
Functional characterisation of catalytically inactive and ISG15-binding mutants of USP18. **(A)** Immunoprecipitation of Flag-tagged wt and mutant forms of USP18 in HEK293T cells co-transfected with wt ISG15 (ISG15.GG) as indicated. Cells were lysed 48 h post-transfection and lysates were immunoprecipitated with Flag-specific antibodies covalently coupled to magnetic beads. Immunoprecipitates (top) and whole cell lysates (WCL; bottom) were subject to immunoblot with antibodies to USP18 and ISG15. **(B)**Immunoprecipitation of V5-tagged IFNAR2 cytoplasmic domain in HEK293T cells co-transfected with STAT2-Myc, USP18.wt-Flag, USP18.I60N-Flag or USP18.C64S-Flag plasmids as indicated. Cells were lysed at 48 h post-transfection and lysates were immunoprecipitated with anti-V5 antibody coupled to protein G dynabeads. following immunoblotting, immunoprecipitates (V5-IP; top) and proteins from whole cell lysates (WCL; bottom) were detected with antibodies to anti-V5 epitope tag, STAT2 and USP18. **(C)** Schematic presentation of the lentiviral technology used to reconstitute USP18 expression in A549-USP18^−/−^ using an inducible system where USP18 expression is driven by the ISG15 promoter (pr15). **(D)** Immunoblot analysis of USP18 expression induced by IFN-α treatment. A549 (CNTL), USP18^−/−^ and USP18.wt-,I60N-,C64S-expressing cell lines were treated with 1000 IU/mL IFN-α for 48 h or left untreated. Whole cell extracts were prepared and USP18, ISG15, MxA and β-actin protein levels were analysed by immunoblot.

Next, we asked whether the USP18.I60N mutant could still be recruited to the IFNAR2 signalling complex despite its inability to interact with ISG15. Previous studies have shown that STAT2 recruits USP18 to the IFNAR2 receptor, where it interferes with the cytosolic interactions between the type I IFN receptor subunits ^14,17,31^. We co-expressed a V5-tagged version of the IFNAR2 cytosolic domain (aa 265-515) and Myc-tagged STAT2 with Flag-tagged wt USP18, USP18.I60N or USP18.C64S in HEK293T cells and performed co-IP assays using anti-V5 antibody coupled to protein G Dynabeads (Fig. 5B). As expected, the IFNAR2 cytoplasmic domain interacted with STAT2 and wtUSP18 (Fig. 5B). Intriguingly, both mutant forms of USP18 co-immunoprecipitated with the IFNAR2 receptor subunit (Fig. 5B), demonstrating that neither the I60N or C64S point mutations disrupted the recruitment of USP18 to the receptor or the subsequent formation of the USP18-dependent type I IFN receptor inhibitory complex.

In order to evaluate the functional consequences of these point mutants, we used CRISPR/Cas9 genome editing to knock-out USP18 in A549 cells followed by lentiviral transduction to reconstitute IFN-inducible expression of Flag-tagged wt or mutant forms of USP18 in the A549-USP18^−/−^ cell line (Fig. 5C). To evaluate USP18 expression, A549 NC1-control cells, which express a negative control guide RNA (NC1) that is nontargeting in humans, A549-USP18^−/−^ and USP18-reconstituted derivatives were treated with IFN-α for 48 h and cell lysates were subjected to immunoblot analysis (Fig. 5D). Interestingly, the expression levels of the reconstituted forms of USP18 were higher compared to NC1 control (Fig. 5D), perhaps because expression of reconstituted USP18 was driven by the ISG15 promoter, which is strongly IFN-responsive ^32^, instead of its native promoter. Consistent with previous observations ^14^, knockout of USP18 increased the levels of ISG15 conjugates, whereas unexpectedly, reconstitution of wt USP18 resulted in a lower level of ISGylation compared to the NC1 control cells. The expression of USP18.I60N mutant was higher compared to the expression levels of wt USP18 and USP18.C64S; however, the accumulation of ISGylated proteins was only marginally increased compared to the cell line expressing wt USP18 (Fig. 5D). Interestingly, protein expression of ISG15 and MxA appeared to be elevated in A549-USP18^−/−^ and the USP18.I60N-expressing cells, signifying higher levels of JAK/STAT signalling (Fig. 5D). As anticipated, expression of the catalytically inactive mutant USP18.C64S resulted in higher levels of ISGylated proteins compared to the cell line reconstituted with wt USP18 (Fig. 5D), and, similar to its mouse counterpart (Ubp43.C61S) ^14^, resulted in lower ISG15 protein expression compared to wt USP18. These observations indicate a dysregulation of the IFN response in an isopeptidase-independent manner (Fig. 5D).

To further evaluate the importance of ISG15-USP18 binding in the regulation of IFN-α signalling, as described in Fig. 1E, we measured the levels of *MxA* and *HERC5* gene expression in these cells (Fig. 6A). Remarkably, *MxA* and *HERC5* expression was significantly higher in USP168.I60N-expressing derivatives compared to NC1 control (averaging around 2.5-fold), denoting that the lack of ISG15-USP18 interaction in these cells led to a dysregulated IFN response, a phenotype similar to A549-USP18^−/−^ cells (Fig. 6A). Notably, ISG expression levels in cells expressing the USP18.C64S mutant were consistently lower than, but not significantly different to, NC1 control cells.

**Figure 6.**
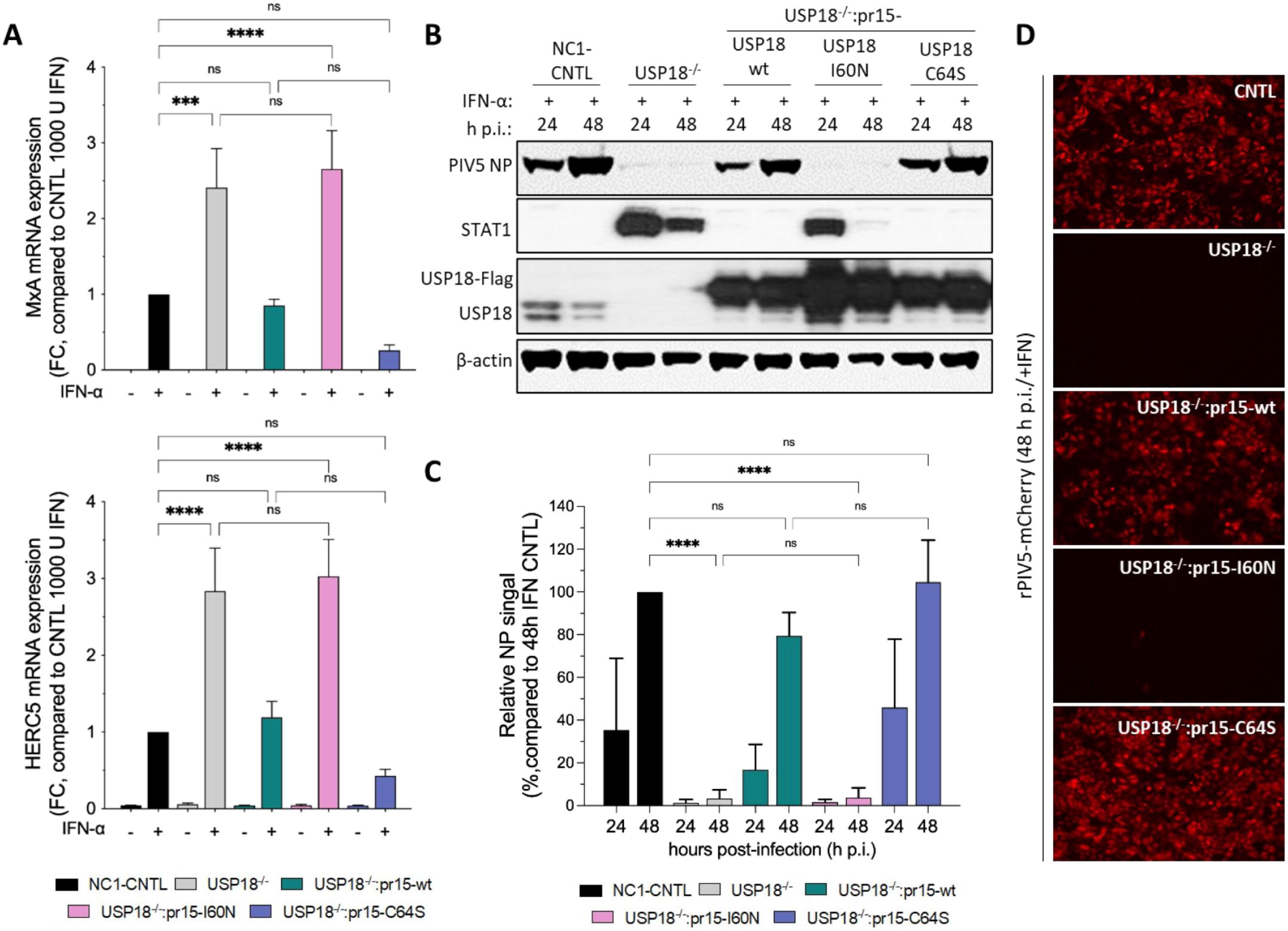
ISG expression is dysregulated in cells expressing USP18 mutant unable to bind ISG15. **(A)** A549 control cells expressing NC1 non targeting guide RNA (NC1-CNTL), A549-USP18^−/−^ and USP18.wt, I60N-, C64S-expressing derivatives (where USP18 expression was under the control of the ISG15 promoter (pr15) and therefore inducible by IFN) were treated with 1000 IU/mL IFN-α for 16 h. Expression of ISGs was tested using reverse transcription quantitative PCR (RT-qPCR) with primers specific for MxA and HERC5. Relative expression was determined following SYBR Green quantitative PCR (qPCR) using ΔΔCt method. β-Actin expression was used to normalize between samples. Data shown represent mean values from three independent experiments performed on different occasions; error bars = SD. Statistical significance was assessed using two-way ANOVA and Tukey multiple comparisons test; *, p < 0.05, **, p < 0.01, ***, p < 0.001, n.s., no statistical significance. **(B)** A549 NC1-CNTL, A549-USP18^−/−^ and USP18.wt-, I60N-, C64S-expressing derivatives were pre-treated with 1000 IU/mL IFN-α for 16 h and then infected with rPIV5-mCherry (MOI 10). Cells were harvested at 24 and 48 h.p.i. and processed for immunoblot analysis using antibodies specific for PIV5 NP, STAT1, USP18 and β-actin. (**C)** Experiments described in (B) were performed independently three times (infections were performed on three separate occasions), and NP and β-actin levels were quantified using Image Studio software (LI-COR Biosciences). Signals were normalised to IFN-α-treated A549 cells infected for 48 h p.i. (set to 100%). Data shown represent mean values from three independent experiments; error bars = SD. Statistical significance was assessed using two-way ANOVA and Tukey multiple comparisons test; ****, p < 0.0001, n.s., no statistical significance. **(D)** Fluorescent imagining of mCherry expression, indicative of rPIV5-mCherry infection, at 48h p.i time point of experiment described in (B).

We predicted that the elevated expression of ISGs in USP18.I60N-expressing cells would engender resistance to infection. Using the assay described in Fig. 2, we observed that IFN-α-pretreated USP18.I60N-expressing cells were largely resistant to infection, as similar to A549-USP18^−/−^ cells, expression of PIV5 NP was barely detectable (Fig. 6B). Quantitative analysis of NP expression levels showed that viral replication was reduced more than 95% in IFN-α-pretreated A549-USP18^−/−^ and USP18.I60N-expressing cells, whereas the levels of viral infection in cell lines expressing the wt and the catalytically inactive mutant USP18.C64S were similar to NC1 control, confirming regulation in these cells (Fig. 6C). mCherry expression levels further verified that IFN-α-pretreatment constrained virus replication in A549-USP18^−/−^ and USP18.I60N-expressing cells (Fig. 6D). Altogether, these experiments demonstrate that disruption of the ISG15-USP18 interaction enhances IFN-mediated signalling, suggesting that ISG15 plays a crucial role in the negative regulation of IFN signalling beyond its indirect function as a USP18 stabiliser.

### The ISG15-USP18 interaction is important for the IFN-α-induced desensitization of IFN-α signalling

Previous studies have shown that USP18 is crucial for the IFN-α-induced desensitization of IFN-α signalling by disrupting the recruitment of IFNAR1 into the ternary IFN-α-IFNAR1-IFNAR2 complex, decreasing the activation of signalling ^16,17,31,33^. To test whether the ISG15-USP18 interaction is important for this USP18-dependent negative regulation of IFN receptor plasticity, we established a desensitization assay based on previous reports ^16^, where A549 NC1-control cells, A549-USP18^−/−^ and USP18-reconstituted derivatives were primed with IFN-α for 8 h or left untreated, washed extensively and then maintained in medium without IFN for 16 h. During the prime-rest phase, USP18 is expressed and prevents further signalling activation following additional stimulation. Following the 16-h resting period, cells were stimulated with IFN-α for 30 min and cell lysates were subjected to immunoblotting to assess phosho-STAT1 expression, which is indicative of early activation of IFN signalling (Fig. 7A). As expected, priming with IFN-α decreased the responsiveness of NC1-control cells to subsequent IFN stimulation as similar levels of phosho-STAT1 were detected in primed and non-primed control cells. A similar phenotype was observed in A549-USP18^−/−^ cells reconstituted with wt USP18 or the catalytically inactive mutant USP18.C64S, indicating that the sensitivity of these cells to IFN-α was downregulated following priming with IFN-α (Fig. 7A). Consistent with previous reports ^16^, A549-USP18^−/−^ cells were not desensitized to IFN-α resulting to 3.8-fold increase in STAT1 phosphorylation when primed with IFN-α compared to the non-primed control (Fig. 7B). Remarkably, cells expressing the USP18.I60N mutant retained their responsiveness toward IFN-α similar to A549-USP18^−/−^ cells (Fig. 7A). Specifically, STAT1 phosphorylation in USP18.I60N-expressing cells showed 3.2-fold increase following IFN-α priming compared to the non-primed control (Fig. 8B). We also performed desensitization assays in our reconstituted A549-ISG15^−/−^ cells (Fig. 8C,D). Here, the degree of desensitization followed the degree of ISG expression (Fig. 1E). Overall, our data strongly suggest that the ISG15-USP18 interaction is important for the USP18-dependent regulation of IFNAR ternary complexes.

**Figure 7.**
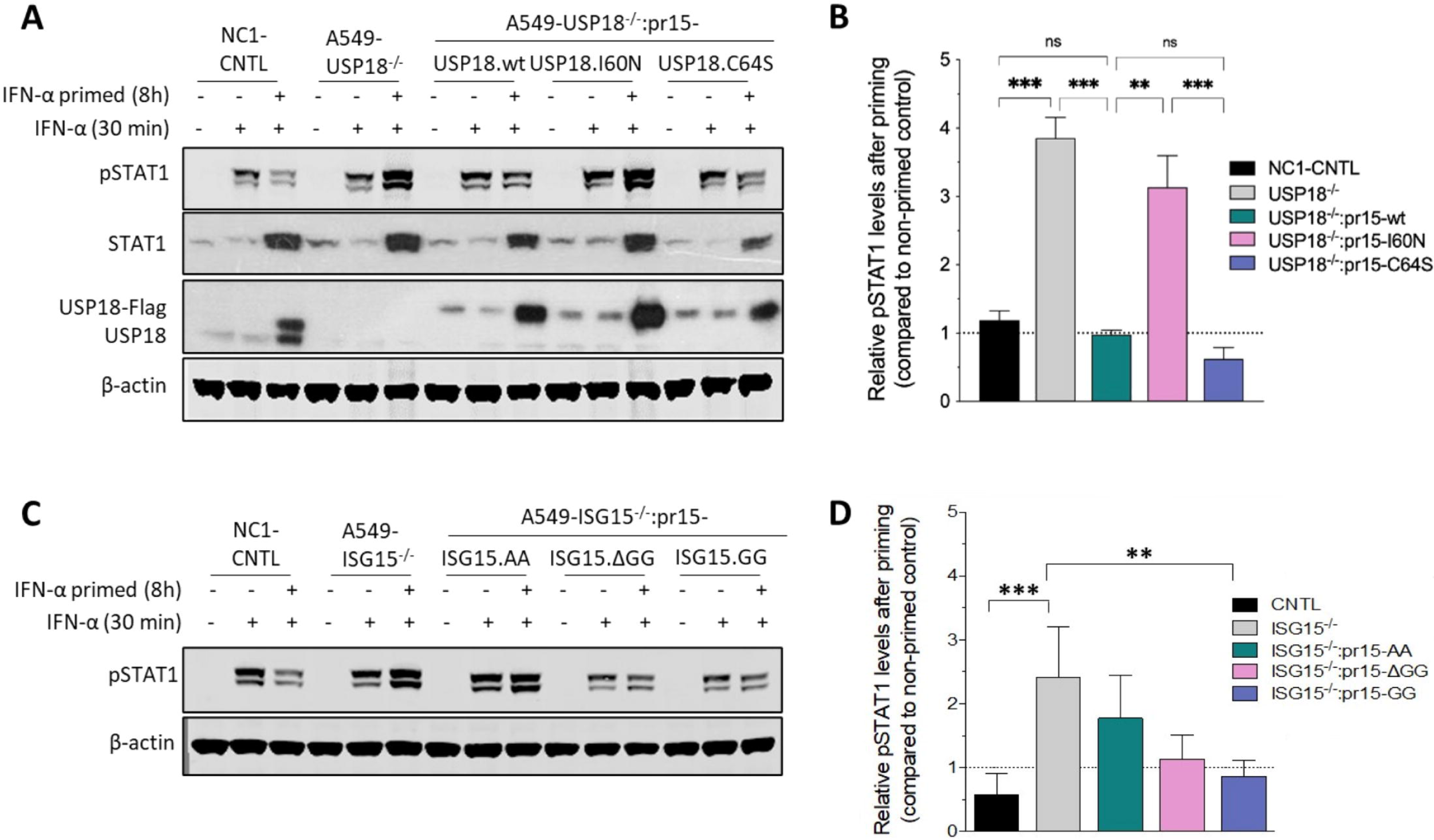
The ISG15-USP18 interaction is important for desensitization of IFN-α signalling. **(A)** A549 control cells expressing NC1 non targeting guide RNA (NC1-CNTL), A549-USP18^−/−^ and USP18.wt-, I60N, C64S-expressing cells were primed with 2000 IU/mL IFN-α (equivalent to 20 ng/mL) for 8 h or left untreated. Cells were washed and re-incubated in medium without IFN for 16 h and then stimulated with 2000 IU/mL IFN-α for 30 min. Cell lysates were subject to immunoblot analysis with antibodies to anti-phospho-STAT1, USP18 and β-actin. Reconstituted USP18 expression was under the control of a clones ISG15 promoter (pr15) and was therefore IFN-inducible. **(B)** Experiments described in (A) were performed independently three times, and phospho-STAT1 and β-actin levels were quantified using Image Studio software (LI-COR Biosciences). Signals are presented as ratios of primed to non-primed control (a ratio of 1 is equivalent to no change). **(C,D)** Experiments in (A) and (B) were repeated with A549 (NC1-CNTL), A549-ISG15^−/−^ and ISG15.AA- or ISG15.GG-expressing cells. Reconstituted ISG15 expression was under the control of a clones ISG15 promoter (pr15) and was therefore IFN-inducible. Data shown represent mean values from three independent experiments; error bars = SD. Statistical significance was assessed using one-way ANOVA and Tukey multiple comparisons test; *, p < 0.05, **, p < 0.01, ***, p < 0.001, ****, p < 0.0001, n.s., no statistical significance.

## Discussion

The negative regulation of the type I IFN system is controlled at multiple levels by a variety of mechanisms, involving sequestration of effector molecules and post-translational modifications, such as ubiquitination or dephosphorylation (reviewed by Arimoto et al. ^34^). Humans with ISG15-deficiency display abnormally strong type I IFN immunity highlighting the emerging role of ISG15 as a central regulator of immunity ^18,19,22,24^. Current models suggest that ISG15 antagonizes the SKP2-mediated ubiquitination and degradation of USP18, promoting its functions ^19,21,24^. We extend these findings and show that the ISG15-dependent stabilisation of USP18 is necessary *but not sufficient* to regulate IFN-I signalling and that non-covalent binding of ISG15 and USP18, via ISG15’s C-terminal tail, is also necessary to facilitate USP18’s inhibitory function. Intriguingly, because this is only – currently – known to occur in human cells (mouse Isg15 is dispensable for Usp18 stability and the regulation of type I IFN signalling), it may suggest this binding mechanism, normally reserved for deISGylation (in all species tested), has been evolutionarily co-opted.

It is well known that the conserved C-terminal di-Gly motif of ISG15 is essential for ISGylation ^35,36^; therefore, the ISGylation-deficient mutants ISG15.AA and ISG15.ΔGG have been extensively used for exploring the functional consequences of ISGylation. Here, we report these ISG15 C-terminal mutants display different propensities for non-covalent binding to USP18 but despite this, both mutants stabilize USP18 comparable to wt ISG15 (Fig. 3). However, when Trp123 in ISG15 was mutated (while maintaining the wt C-terminal tail), it was unable to stabilise USP18, showing that interactions are likely required. To achieve high affinity binding, the ISG15 C-terminal tail engages with the catalytic pocket of USP18 via a two-step binding process; initial hydrophobic interactions centred on ISG15 Trp123 (Trp121 in mouse, Fig. 4A) and the ISG15 binding box 1 (IBB-1) in USP18 induce rearrangement of the USP18 ‘switching loop’ thus allowing the ISG15 C-terminal tail access to the catalytic site in USP18 ^12^. Indeed, Basters and colleagues (2017)^12^ showed that, *in vitro*, Isg15-W121R:Usp18 interactions were virtually abolished and that the C-terminal tail of ISG15 contributes the most affinity. Interaction studies with ISG15-AA and -ΔGG mutants showed that the C-terminal tail of ISG15 is important for tight binding, and with ISG15-AA, we could not detect any interaction with USP18, even though Trp123 was present (Fig. 4D). Therefore, the hydrophobic interactions are probably weak or fast dissociation kinetics make them difficult to detect in our assays. It is possible that ISG15-dependent stability of USP18 is concentration dependent particularly if the kinetics of the initial hydrophobic interactions are fast; in that regard, ISG15 is known to be one of the most highly expressed ISGs during the antiviral response, yet USP18 expression is considered low. While these data do not rule out the possibility that indirect interactions between ISG15 (via Trp123) and an unknown partner are responsible for USP18 stability, which may be consistent with previous reports showing that ISG15 can abrogate the USP18-SKP2 complex and rescue USP18 from proteasomal degradation independently of its ability to bind USP18 ^24^.

Importantly, our work suggests that stronger ISG15-USP18 binding (reliant on the ISG15 C-terminal tail) is required for USP18-dependent regulation of the type I IFN response as the level of binding to USP18 faithfully reflected the level of IFN-α signalling regulation and the permissiveness of cells to viral infection. We independently confirmed the importance of a ISG15-USP18 interaction as expression of mutant USP18 unable to bind ISG15 (USP18-I60N) could not regulate type I IFN signalling even though it could interact with the IFNAR2 signalling complex (Fig. 5). We did not observe reduced USP18 abundance in USP18-I60N-expressing cells, suggesting any loss in regulation was not due to lack of USP18 stability. In line with findings using mouse Usp18 ^13,14^, the protease activity of human USP18 remains dispensable for signalling regulation in humans.

Previous studies have shown that USP18 desensitises type I IFN signalling ^16,17,37^. Specifically, STAT2 recruits USP18 to the IFNAR2, where it interferes with the cytosolic interactions between receptor subunits, impeding the recruitment of IFNAR1 into the ternary complex ^17,37^. Remarkably, we demonstrate here that the non-covalent binding of ISG15 and USP18 is required for the USP18-dependent negative regulation of IFNAR dimerization (Fig. 7). Consistent with previous reports ^37^, our data show that ISG15 appears dispensable for the interaction of USP18 IFNAR2; therefore, we hypothesize that the ISG15’s contribution may be essential for further stabilizing the USP18-inhibitory complex, for recruiting yet-to-be identified interaction partners of USP18 or promoting a conformational change in USP18 necessary for its inhibitory activity.

Structural studies have shown that the ISG15-bound USP18 adopts a different conformation, where the ‘switching loop’ in the thumb domain of USP18 acquires an active conformation, enabling access of the LRLRGG C-terminal tail of ISG15 into the catalytic cleft ^12,38^. Here, we showed that replacing the C-terminal di-Gly of ISG15 with di-Ala completely abolishes ISG15-USP18 interaction (Fig. 4). Although these amino acid substitutions are considered subtle, it is possible that the two extra methyl groups present in the di-Ala motif render it inaccessible to the tight catalytic cleft of USP18, abolishing the interaction with the ISG15.AA mutant. This may also explain why the ISG15.ΔGG mutant, which retains the LRLR motif of the C-terminal tail, interacted to a greater extent with USP18 compared to ISG15.AA. Moreover, in line with previous findings ^24^, we showed that the USP18.I60N mutant was unable to bind ISG15 (Fig. 6A). Interestingly, Ile60 does not belong to the ISG15-binding boxes ^12^ but its proximity to the catalytic cysteine (C64) may affect the conformational dynamics of the catalytic cleft, diminishing the interaction with ISG15. Consistent with previous reports (23), we have observed that the catalytically inactive mutant (USP18.C64S) has stronger binding affinity for ISG15, compared to wt USP18, and there is a consistent pattern of stronger negative regulation (lower ISG expression, increased desensitization to IFN-α) in cells expressing this mutant, supporting our observation that the level of binding of ISG15 toward USP18 determines the level of IFN-α signalling regulation. Hence, it is possible that the binding of ISG15 locks USP18 into a more stable structural conformation that may serve its regulatory functions at the level of IFNAR assembly. A caveat to this model is that Isg15 is dispensable for Usp18-mediated regulation of type I IFN signalling in mice ^21^.

That Isg15 does not stabilise Usp18 in mice ^21^, illustrates interesting interspecies variation with regard to ISG15 function. Unlike the highly conserved ubiquitin ^39^, ISG15 possesses remarkable sequence variation between species, with amino acid identity of 65.6% (mature protein) between human and mouse, suggesting that the different biochemical properties of ISG15 between species may be key determinants of ISG15’s species-specific functions ^40,41^. Indeed, it has been shown that human ISG15 associates with higher affinity to USP18 compared to its mouse counterpart ^21^, which in agreement with our findings, suggests that gain-of-function mutations in ISG15 and/or USP18 that facilitate stronger ISG15-USP18 interactions have been evolutionary selected in humans (though it is also possible there has been a loss-of-function in mice). Why this trait is apparent in humans and not mice is of interest and may suggest that an additional, yet-to-be identified factor is taking the place of ISG15 in mice or that humans may require a different level of IFN-regulatory control.

Intriguingly, our proteomics profiling and pathway analyses highlighted the enrichment of components involved in IFN-γ signalling in A549-ISG15^−/−^ and ISG15.AA-expressing cells (Fig. 2). This may be due to the accumulation of STAT1 homodimers as a by-product of enhanced type I immunity, leading to enrichment of GAS-containing genes ^27,28,42^, such as IRF1 and MHC Class I proteins in these cells. IRF1 transcription factor itself is involved in the regulation of genes implicated in antiproliferative ^43^ and antigen processing pathways ^44–46^. These findings support our previous work that demonstrates that ISG15 deficiency leads to translational regression following IFN-α treatment ^22^ and further suggest that intervention strategies that target the ISG15-USP18 interaction may be of therapeutic use in anticancer therapy.

In conclusion, we have demonstrated that intracellular ISG15 is essential to negatively regulate IFN-α responses via its non-covalent interaction with USP18, thereby averting autoinflammatory consequences of uncontrolled type I IFN signalling. This hitherto human-specific trait has possibly been acquired through co-option of a binding mechanism normally reserved for the process of deISGylation. Further investigation is needed to decipher the biophysical properties of the USP18-mediated inhibitory complex at the level of IFNAR assembly and resolve how the binding of ISG15 to USP18 facilitates those interactions.

## Methods

### Cells

h-Tert-immortalized dermal fibroblasts from control (C1) and ISG15-deficient patients (P1) ^18,19^, HEK293T (human embryonic kidney cell), A549 cells (human adenocarcinoma alveolar basal epithelial cells), and derivatives, were were grown as monolayers in Dulbecco’s modified Eagles’s medium (DMEM; Sigma) supplemented with 10% (v/v) foetal bovine serum (FBS, Biowest) and incubated in 5% (v/v) CO_2_ at 37°C in a humidified incubator incubator (C1 and P1 media additionally contained 1x GlutaMAX supplement (ThermoFisher Scientific). A549-ISG15^−/−^ cells (clone B8) were generated as previously described using CRISPR/Cas9n system; transfectants were enriched by treating cells with puromycin (11μg/mL) for 2 days, then single-cell cloned and successful knockout cells were validated by immunoblot analysis ^22^. Lentiviral technology was used to reconstitute expression of wt ISG15.GG (NCBI Ref Seq, NM_005101.3) or C-terminal mutants, ISG15.AA and ISG15.ΔGG in A549-ISG15^−/−^ to generate the following derivative cell lines; A549-ISG15^−/−^:prI5-GG, A549-ISG15^−/−^:prI5-AA, A549-ISG15^−/−^:prI5-ΔGG, respectively. Expression of wt and C-terminal mutant forms of ISG15 was driven by the native ISG15 promoter (pr15) (NCBI Ref seq, NG_033033.2) cloned using our lentiviral vector. An internal ribosome entry site (ISRE) downstream the ISG15 gene allows expression of puromycin resistance (pac) gene following induction of pr15. Hence, for puromycin selections, cells were primed with 1000 IU/mL IFN-α for 4 h prior to treatment with puromycin (11μg/mL) for 2 days. To generate the A549-ISG15^−/−^:UBA7^−/−^:pr15-GG cell line, A549-ISG15^−/−^:pr15-GG cells were further modified to stably express *Streptococcus pyogenes* Cas9 and then transduced with UBA7 sgRNA-expressing lentiGuide-Puro (sgRNA sequences; sense: caccGCACACGGGTGACATCACTG; antisense: aaacCAGTGATGTCACCCGTGTGC) as described previously ^22^. For generating A549-USP18^−/−^ cell lines, A549 cells were modified to stably express *S. pyogenes* Cas9 and then transduced with USP18 sgRNA-expressing lentiGuide-Puro (sgRNA sequences; sense: caccgGGGGCCGCACTGCTTTCTGC; antisense: aaacGCAGAAAGCAGTGCGGCCCCc). Blasticidin/puromycin-resistant A549-USP18^−/−^ cells were single-cell cloned in 96-well plates and successful knockout cells were validated by immunoblot analysis. Lentiviral technology using our pr15 lentiviral vector was used to reconstitute expression of wt USP18 (NCBI Ref seq NM_017414.4) or the point mutants USP18.I60N and USP18.C64S (generated by site-directed mutagenesis) in A549-USP18^−/−^ to generate the A549-USP18^−/−^:prI5-USP18.wt, A549-USP18^−/−^:prI5-USP18.I60N and A549-USP18^−/−^:prI5-USP18.C64S, respectively. All lentiviruses used in this study were generated in HEK293T cells using a previously described self-inactivating lentiviral constitutive expression system ^47^.

### Reverse transcription quantitative PCR

Total RNA was isolated from A549 cells treated with 1000 IU/ml IFN-α for 18 h or from hTert-immortalized dermal fibroblasts treated with 1000 IU/ml IFN-α for 12 h, washed with PBS then incubated in media without IFN-α for a further 24 h. Total RNA was isolated with TRIzol (ThermoFisher Scientific) and Direct-zol RNA Miniprep Plus kits (Zymo Research), including removal of contaminating DNA following DNase I treatment, following the manufacturer’s instructions. A total of 500 ng of RNA was reverse transcribed using LunaScript reverse transcriptase (New England Biolabs) according to the manufacturer’s recommendations. Quantitative PCRs (qPCRs) were performed with PerfeCTa SYBR green SuperMix (Quanta BioScience) using fast two-step cycling performed in a Mx3005P real time thermal cycler (Stratagene) and included an initial 2 min enzyme activation step at 95°C, followed by 40 cycles of 5 s at 95°C and 20 s at 60°C. Melting curve analysis was performed to verify amplicon specificity. For each assay, quantification of *β-ACTIN* mRNA was used to normalize between samples and the average cycle threshold (CT) was determined from three independent cDNA samples from independent cultures (technical replicates). Fold changes were determined from three independent assays performed at different times (biological replicates). Relative expression compared to non-treated control cells was calculated using the ΔΔCT method. Primer sequences were: *MxA* 5’GCCTGCTGACATTGGGTATAA and 5’CCCTGAAATATGGGTGGTTCTC, *HERC5* 5’GACGAACTCTTGCACCGTCTC and 5’GCGTCCACAGTCATTTTCCAC, *β-ACTIN* 5’AGCGAGCATCCCCCAAAGTT and 5’AGGGCACGAAGGCTCATCATT.

### Viral infections and IFN treatment

Recombinant mCherry-expressing parainfluenza virus type 5 (rPIV5-mCherry) (provided by Professor Biao He, University of Georgia) ^48^ stocks were prepared by inoculating Vero cells at a multiplicity of infection (MOI) of 0.001 with continual rocking at 37°C. Supernatants were harvested at 2 d p.i., clarified by centrifugation at 3,000 x g for 15 min, aliquoted and snap frozen. Titres were estimated by standard plaque assay on Vero cells in 6-well plates.

For virus resistance assays, cell monolayers at 70-80% confluency (A549) or full confluency (hTert-immortalized fibroblasts) were infected in 6-well plates with virus diluted in 1 mL media supplemented with 2% (v/v) FBS to achieve an MOI of 10. Virus adsorption was for 1 h with continual rocking at 37°C, after which 1 mL of media supplemented with 2% (v/v) FBS was added to the viral inoculum and incubated in 5% (v/v) CO_2_ at 37°C until harvested. When cells were treated with IFN-α prior to infection (pre-treated) this was done with 1000 IU/mL IFN-α2b (referred to as IFN-α from here on; IntronA, Merck Sharp & Dohme Ltd). IFN-α remained on cells for the duration of experiments. Cells were either processed for immunoblot analysis or observed with fluorescence microscopy using EVOS M5000 Imaging System (pictures taken at 10X magnification).

### Immunoblotting

Confluent monolayers in 6-well dishes were lysed with 250 µL (A549s) or 100 µL (hTert-immortalized dermal fibroblasts) 2 x LaemmLi sample buffer (4% w/v SDS, 20% v/v glycerol, 0.004% w/v bromophenol blue and 0.125 M Tris-HCl, pH 6.8 with 10% v/v β-mercaptoethanol) for 10 min, incubated at 95°C for 10 min, sonicated at 4°C with 3 cycles of 30 s on 30 s off in a Bioruptor Pico (Diagenode) and clarified by centrifugation at 12,000 x g, 4°C for 10 min. SDS-PAGE in Tris-glycine-SDS running buffer and immunoblotting followed standard techniques using the following antibodies: mouse monoclonal anti-ISG15 F-9 (Santa Cruz Biotechnology Cat# sc166755), rabbit polyclonal anti-ISG15 H-150 (Santa Cruz Biotechnology Cat# sc50366), rabbit polyclonal anti-MxA (Proteintech Cat# 13750-1-AP), mouse monoclonal anti-UBA7 (anti-UBE1L B-7; Santa Cruz Biotechnology Cat# sc-390097), rabbit anti-USP18 (Cell Signalling Technology Cat# 4813S), mouse monoclonal anti-STAT1 (N-terminus; BD Transduction Laboratories Cat# 610116), rabbit monoclonal anti-phosphorylated STAT1 (anti-phospho-STAT1 (Tyr701); Cell Signalling Technology Cat# 9167), mouse monoclonal anti-PIV5 NP 125 ^49^, mouse anti-V5 tag Pk 336 ^49^, antibody mouse monoclonal anti-β-actin (Sigma Cat# A2066). For quantitative immunoblots primary antibody-probed membranes were incubated with IRDye secondary antibodies (LiCOR) and signals detected using an Odyssey CLx scanner. Data were processed and analysed using Image Studio software (LiCOR).

### Immunoprecipitation

Prior to immunoprecipitation, HEK293T cells were transiently transfected to express the proteins of interest using calcium-phosphate coprecipitation transfections. One day prior to transfection, HEK293T cells were seeded in 6-well plates such that they were logarithmically growing on the day of transfection (i.e., 50-60% confluent). A total of 200 μL calcium-phosphate precipitate was prepared for each well by mixing each plasmid DNA (diluted in total 90 μL in dΗ_2_Ο) with 10 μL of 2.5M CaCl_2_ solution. DNA/CaCl_2_ solutions were added dropwise into 100 μL of 2× HEPES-buffered saline (HeBS) (50 mM HEPES, 0.28 M NaCl, 10 mM KCl, 1.5 mM Na_2_HPO_4_, 12 mM D-glucose, pH 7.05) and incubated for 20 min at room temperature (RT). Chloroquine diphosphate solution was added to cell culture media to 25 µM final concentration and calcium phosphate precipitate was added dropwise onto plate and mixed gently. At 16 h after transfection, the calcium phosphate precipitate was removed, and cells were incubated for further 24 h before co-immunoprecipitation (co-IP) assays. The following vectors were used for co-IP assays: pLHCX-STAT2-Myc (kind gift from Dr Michael Nevels, St Andrews University), pcDNA3.1-USP18.wt-3XFlag, pcDNA3.1-USP18.I60N-3XFlag, pcDNA3.1-USP18.C64S-3XFlag, pcDNA3.1.ISG15.GG and pcDNA3.1-IFNAR2.CTD-V5, which expresses the cytoplasmic C-terminal domain (CTD) of IFNAR2.

For co-IP assays, confluent monolayers of IFN-treated A549 derivatives grown in T150 cm^2^ flasks or plasmid-transfected HEK293T cell cultures grown in 6-well plates were harvested in phosphate-buffered saline (PBS), pelleted by centrifugation (300 × g, 5 min, 4°C) and resuspended in 1 mL co-IP lysis buffer (50 mM Tris pH 7.5, 150 mM NaCl, 0.1% [v/v] Triton-X) supplemented with 1x cOmplete^TM^ protease inhibitor cocktail (Merck). The supernatant was separated by centrifugation (12,000 x g, 10 min, 4°C) and incubated overnight at 4°C with gentle rotation with 40 μL Pierce Anti-c-Myc Magnetic Beads (ThermoFisher Scientific Cat# 88842) or anti-Flag® M2 Magnetic Beads (Merck Cat# M8823) or anti-V5 tag antibody coupled with Protein G Dynabeads^TM^ (Invitrogen) by following manufacturer’s instructions. Complexes were washed three times with co-IP wash buffer (1X TBS; 50 mM Tris pH 7.5, 150 mM NaCl), incubated in 50 μL 1X NuPage LDS sample buffer (ThermoFisher Scientific) for 20 min at RT and then, further incubated for 10 min at 70°C. Beads were magnetically separated and 10% (v/v) of β-mercaptoethanol was added in the eluted supernatant containing target antigens. Immunoprecipitates were subjected to SDS-PAGE and immunoblotting.

### IFN signalling Desensitization assay

Cells grown to 70-80% confluency in 6-well plates were primed with 2000 IU/mL IFN-α (equivalent to 20 ng/mL) for 8 h or left untreated. Following priming, cells were washed four times with PBS and maintained in medium without IFN for 16 h (resting period) and then stimulated with 2000 IU/mL IFN-α for 30 min. Cell lysates were subjected to SDS-PAGE and immunoblotting.

### Tandem mass tags (TMT)-based quantitative proteomics

Tandem mass tags (TMT)-based proteomics was used to quantify differences in protein abundance following IFN-α treatment in A549 control cells, A549-ISG15^−/−^ and the A549-ISG15^−/−^:prI5-AA and A549-ISG15^−/−^:prI5-GG derivatives. Cells grown to 60-70% confluency in T25 flasks were treated with 1000 IU/mL IFN-α for 24, 48 and 72 h or left untreated for 72h. Whole cell lysate protein digestion was performed as described before ^50^ for each time point. For lysis, cells were washed twice with PBS, and 250 mL lysis buffer added (6 M Guanidine/50 mM HEPES pH 8.5). Cells were scraped in lysis buffer, vortexed extensively and then sonicated. Cell debris was removed by centrifuging at 21,000 g for 10 min, twice. For the Dithiothreitol (DTT) was added to a final concentration of 5 mM and samples were incubated for 20 min. Cysteines were alkylated with 14 mM iodoacetamide and incubated 20 min at room temperature in the dark. Excess iodoacetamide was quenched with DTT for 15 min. Samples were diluted with 200 mM HEPES pH 8.5 to 1.5 M Guanidine followed by digestion at room temperature for 3 h with LysC protease at a 1:100 protease-to-protein ratio. Samples were further diluted with 200 mM HEPES pH 8.5 to 0.5 M Guanidine. Trypsin was then added at a 1:100 protease-to-protein ratio followed by overnight incubation at 37°C. The reaction was quenched with 5% formic acid, then centrifuged at 21,000 g for 10 min to remove undigested protein. Peptides were subjected to C18 solid-phase extraction (SPE, Sep-Pak, Waters) and vacuum-centrifuged to near-dryness.

Samples were prepared for TMT labelling as previously described ^50^. Desalted peptides were dissolved in 200 mM HEPES pH 8.5 and 25 mg of peptide labelled with TMT reagent. TMT reagents (0.8 mg) were dissolved in 43 mL anhydrous aceto-nitrile and 3 mL added to peptide at a final acetonitrile concentration of 30% (v/v). Following incubation at room temperature for 1 h, the reaction was quenched with hydroxylamine to a final concentration of 0.3% (v/v). Sample labelling was performed using 16-plex labelling reagent (ThermoFisher Scientific^TM^ CAT# A44520) and TMT-labelled samples were combined at a 1:1:1:1:1:1:1:1:1:1:1:1:1:1:1:1 ratio. Samples were vacuum-centrifuged to near dryness and subjected to C18 SPE (Sep-Pak, Waters). An unfractionated single shot was analysed initially to ensure similar peptide loading across each TMT channel, thus avoiding the need for excessive electronic normalization. As all normalisation factors were >0.5 and <2, data for each singleshot experiment was analysed with data for the corresponding fractions to increase the overall number of peptides quantified.

TMT-labelled tryptic peptides were subjected to pH reversed-phase (HpRP) fractionation using an Ultimate 3000 RSLC UHPLC system (Thermo Fisher Scientific) equipped with a 2.1 mm internal diameter (ID) x 25 cm long, 1.7 mm particle Kinetix Evo C18 column (Phenomenex). Mass spectrometry data was acquired using an Orbitrap Lumos and an ultimate 3000 RSLC nano UHPLC equipped with a 300 mm ID x 5 mm Acclaim PepMap m-Precolumn (Thermo Fisher Scientific) and a 75 mm ID x 50 cm 2.1 mm particle Acclaim PepMap RSLC analytical column was used as described before ^50^.

### Data analysis of MS spectra

For MS3-based TMT, as previously described ^50^, TMT tags on lysine residues and peptide N termini (229.162932 Da) and carbamidomethylation of cysteine residues (57.02146 Da) were included as static modifications. Proteins were quantified by summing TMT reporter ion counts across all matching peptide-spectral matches using ‘MassPike’, as described previously ^51^. Briefly, a 0.003 Th window around the theoretical m/z of each reporter ion (126, 127 n, 128 n) was scanned for ions, and the maximum intensity nearest to the theoretical m/z was used. An isolation specificity filter with a cutoff of 50% was employed to minimise peptide co-isolation ^51^. Peptide-spectral matches with poor quality MS3 spectra (more than 3 TMT channels missing and/or a combined S:N ratio of less than 100 across all TMT reporter ions) or no MS3 spectra at all were excluded from quantitation. Peptides meeting the stated criteria for reliable quantitation were then summed by parent protein, in effect weighting the contributions of individual peptides to the total protein signal based on their individual TMT reporter ion yields. Protein quantitation values were exported for further analysis in Excel.

For protein quantitation, reverse and contaminant proteins were removed, then each reporter ion channel was summed across all quantified proteins and normalised assuming equal protein loading across all channels. Protein hits quantified by a single peptide were removed from the dataset. The expression profile of each protein was observed after comparing protein abundance to the condition (cell line/time point) with the highest MS intensity score (set to 1) and normalised values were plotted against each time point for each cell line (see Plotter in Supplementary File 1)

### Pathway Analysis

To identify individual ISGs from our dataset, a list of 7112 gene symbols were searched in ‘Interferome v2.01’ (http://interferome.its.monash.edu.au/interferome/home.jspx) ^26^. The Interferome analysis was conducted on fold change values, which were calculated by dividing the MS intensity score of each identified protein at a given time point post IFN treatment (24, 48, 72 h) by the MS intensity score of the untreated control for each cell line. A protein hit was considered to be an ISG if it was upregulated at least 1.7-fold in A549 control cells following IFN-α treatment.

The Database for Annotation, Visualization and Integrated Discovery (DAVID) version 6.8 (https://david.ncifcrf.gov) was used to stringently identify cell line-specific enriched pathways. An ‘enrichment ratio’ for each protein was obtained for each time point as follows (MIS is MS intensity score):

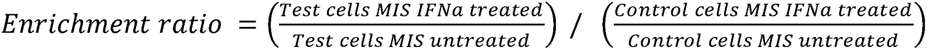

Protein hits upregulated at least 1.7-fold following enrichment were submitted using the UniProt accession number and default medium classification stringency. Clusters were considered significant if the Benjamini-Hochberg adjusted p-value was <0.05.

## Data availability

The mass spectrometry proteomics data have been deposited to the ProteomeXchange Consortium (http://www.proteomexchange.org) via the PRIDE ^52^ partner repository with the dataset identifier (xxxx).

## Acknowledgments

In memory of Dr Christina Paulus. We are indebted to technical assistance provided by Dan Young. This work was supported by grants from the Academy of Medical Sciences (SBF003/1028 to DJH), Wellcome Trust Institutional Strategic Support Fund (to DJH), Wellcome Trust (101788/Z/13/Z to RER), the UK Medical Research Council (MC_UU_12014/1 to JM and CGGB) and by the Wellcome Trust via a Senior Clinical Research Fellowship (108070/Z/15/Z to MPW).

