## Supplementary figures and images for "A co-opted ISG15-USP18 binding mechanism normally reserved for deISGylation controls type I IFN signalling"

### Supplemental Figure 1

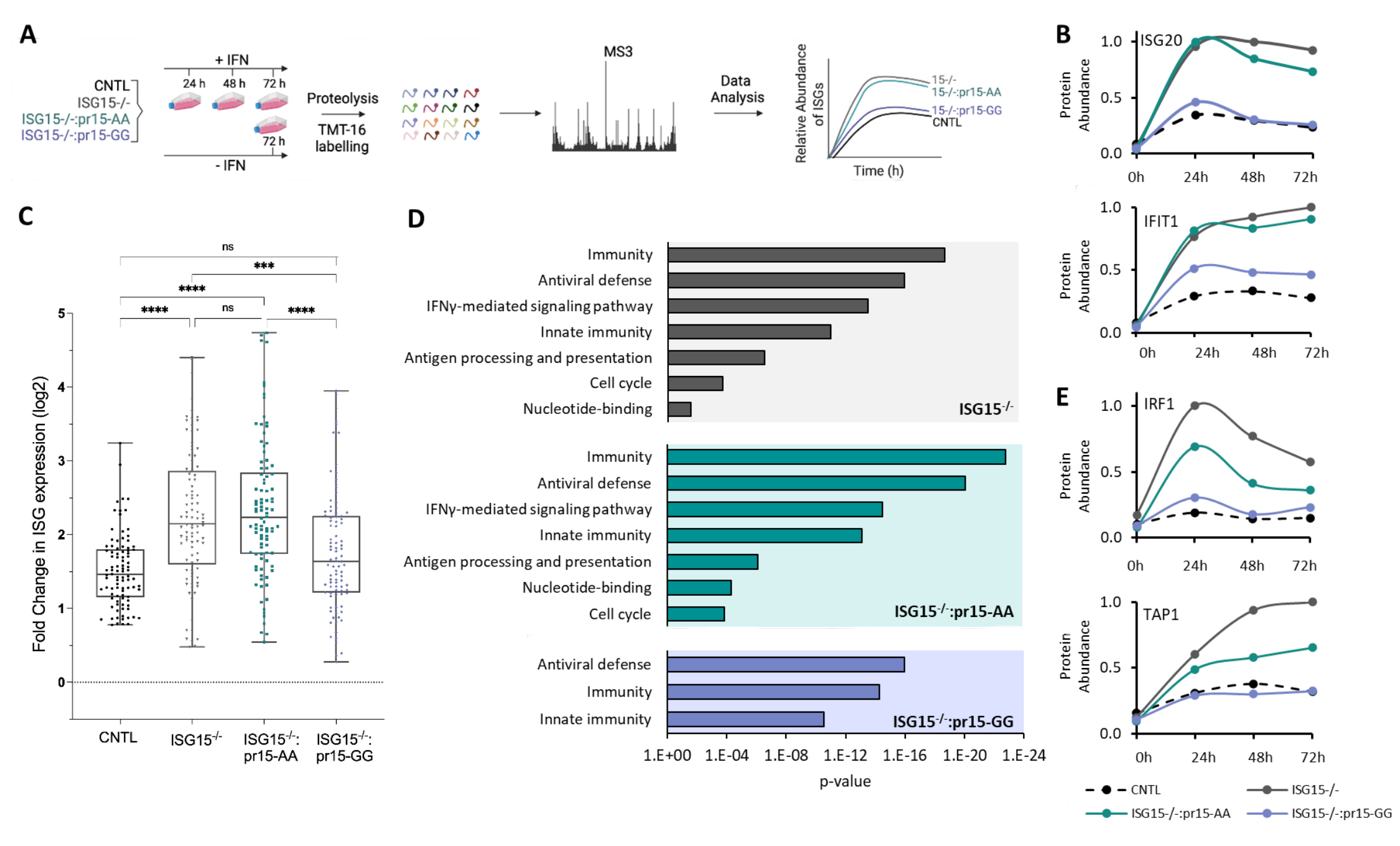

### Supplemental Figure 2

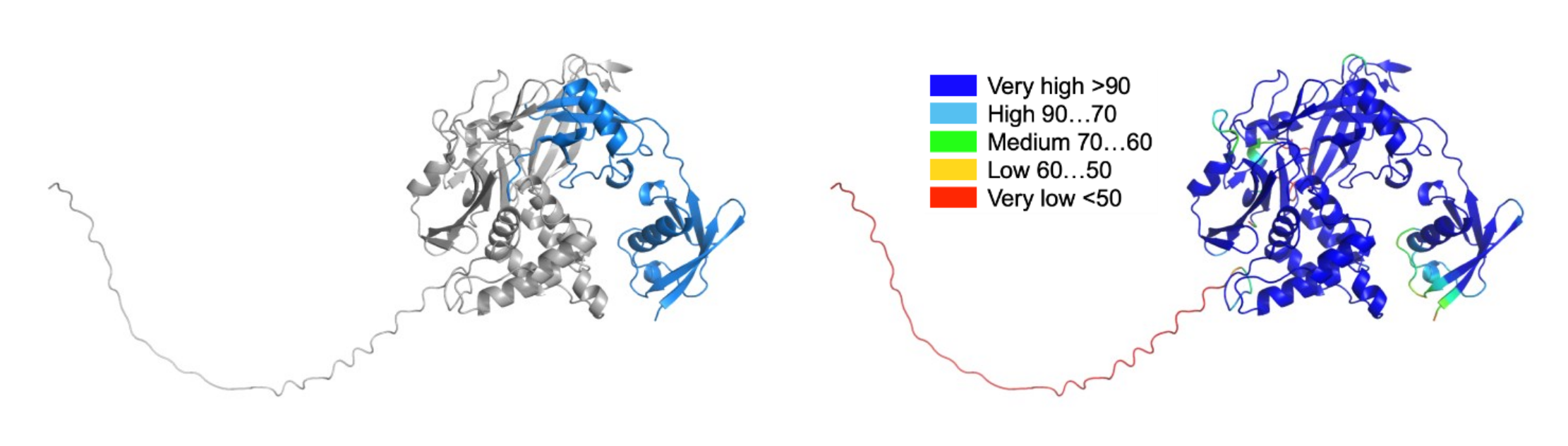
